# Mosaic Evolution of Beta Barrel Porin Encoding Genes in *Escherichia coli*

**DOI:** 10.1101/2021.09.21.461324

**Authors:** Xiongbin Chen, Xuxia Cai, Zewei Chen, Jinjin Wu, Gaofeng Hao, Quan Luo, Guoqiang Zhu, Wolfgang Koester, Aaron P White, Yi Cai, Yejun Wang

**Affiliations:** Youth Innovation Team of Medical Bioinformatics, Shenzhen University Health Science Center, Shenzhen 518060, China; Department of Stomatology, Shenzhen Maternal and Child Health Care Hospital Affiliated to Southern Medical University, China; College of Veterinary Medicine, Yangzhou University, Yangzhou, China; Vaccine and Infectious Disease Organization-International Vaccine Centre, Saskatoon, Saskatchewan, Canada; Department of Medical Microbiology, School of Basic Medicine, Shenzhen University Health Science Center, Shenzhen, China

**Keywords:** Mosaic Evolution, local recombination, β-barrel porin, FhuA, *Escherichia coli*, LamB, OmpA, OmpC, OmpF

## Abstract

Bacterial porins serve as the interface interacting with extracellular environment, and are often found under positive selection to fit in different environmental stresses. Local recombination has been identified in a handful of porin genes to facilitate the rapid adaptation of bacterial cells. It remains unknown whether it is a common evolutionary mechanism in gram-negative bacteria for all or a majority of the outer membrane proteins. In this research, we investigated the β-barrel porin encoding genes in *Escherichia coli* that were reported under positive Darwinia selection. Besides *fhuA* that was found with ingenic local recombination predominantly previously, we identified four other genes, i.e., *lamB, ompA, ompC* and *ompF*, all showing the similar mosaic evolution patterns as in *fhuA*. Comparative analysis of the protein sequences disclosed a list of highly variable regions in each protein family, which are mostly located in the convex of extracellular loops and coinciding with the binding sites of various bacteriophages. For each of the porin family, mosaic recombination leads to various combinations of the HVRs with different sequence patterns, generating diverse protein groups. Structure modeling further indicated the conserved global topology for various groups of each porin family, but the extracellular surface varies a lot that is formed by individual or combinatorial HVRs. The conservation of global tertiary structure ensures the channel activity while the wide diversity of HVRs may assist bacteria avoiding the invasion of phages, antibiotics or immune surveillance factors. In summary, the study identified multiple bacterial porin genes with mosaic evolution, a likely general strategy, by which outer membrane proteins could facilitate the host bacteria to both maintain normal life processes and evade the attack of unflavored environmental factors rapidly.

**Importance:** Microevolution studies can disclose more elaborate evolutionary mechanisms of genes, appearing especially important for genes with multifaceted function such as those encoding outer membrane proteins. However, in most cases, the gene is considered as a whole unit and the evolutionary patterns are disclosed. In this research, we reported that multiple bacterial porin proteins follow mosaic evolution, with local ingenic recombination combined with spontaneous mutations based positive Darwinia selection, and conservation for most of the other regions. It could represent a common mechanism for bacterial outer membrane proteins. The study also provides insights on development of new anti-bacterial agent or vaccines.

## Introduction

β-barrel outer membrane proteins (OMPs) exert multiple activities in gram-negative bacteria, for instances, transport of various substrates and signal transduction, facilitating bacterial fitting in different environment (Rollauer et al., 2015). Typically, the β-barrel OMPs form closed barrels embraced by anti-parallel β-strands with hydrophobic residues toward bacterial outer membrane and hydrophilic residues toward the inner surface of the channels (Rollauer et al., 2015; Kleinschmidt 2015). Multiple β-barrel OMP encoding genes were found under positive Darwinia selection in *Escherichia coli*, including *fhuA, ompA, ompC, ompF*, etc (Chen et al., 2006; Petersen et al., 2007). However, recently, one of the genes *fhuA* was reported with local mosaic recombination rather than positive Darwinia selection mediated by spontaneous mutations (Wang et al., 2018). Similarly, local recombination was also reported in *ompA* and *ompF* in *Chlamydia* and *Yersinia* respectively (Millman et al., 2001; Stenkova et al., 2011). It remains unknown whether there are other *E. coli* β-barrel OMP genes also showing mosaic evolution patterns as in *fhuA*.

Like FhuA, many other β-barrel OMPs reported under positive selection are also receptors for various substrates and have multifaceted functions, e.g., OmpA, OmpC and OmpF. OmpA is a major heat-modifiable porin with a molecular weight of 28-36 kDa. The N-terminus of OmpA forms an eight-stranded, anti-parallel, hydrophobic β-barrel domain while the C-terminus forms a hydrophilic, globular domain located in the periplasm (Lopez-Barbosa et al., 2020). OmpA participates in various processes of bacterial infections, such as bacterial adhesion, invasion, intracellular survival, and escape from host defenses (Confer and Ayalew 2013; Krishnan and Prasadarao 2012). OmpA is also the receptor of multiple phages and colicins (U and L) and is good candidates for vaccine design (Krishnan and Prasadarao 2012; Pore and Chakrabarti 2013; Futse et al., 2019). OmpC and OmpF are classical porin proteins, abundantly distributed in *E. coli* outer membrane and essential for bacterial survival in extremely acidic conditions (Bekhit et al., 2011). They are also the main channels via which antibiotics and other virulence factors enter bacterial cells (Misra and Benson 1988; Yoshimura and Nikaido 1985; Bafna et al., 2020). The decreased expression of these genes could be related with bacterial resistance to the antibiotics (Chetri et al., 2019). Like OmpA and FhuA, OmpC and OmpF are also the receptors for bacteriophages (Traurig and Misra 1999; Parent et al., 2014; Chen et al., 2020).

In this study, we investigated the β-barrel OMP genes under positive selection reported previously including *ompA, ompC, ompF* and others, and observed whether they showed mosaic evolution patterns with local recombination as in *fhuA*. Furthermore, sequence analysis and structure modeling were performed to the positive genes, to explore the underlying mechanisms and functional relevance of the mosaic evolution.

## Results

### 1. Screening of *Escherichia* porin genes under mosaic evolution

Besides FhuA, the protein sequences of four other beta barrel porins (LamB, OmpA, OmpC and OmpF) also showed the mosaic-clustering patterns, different from the phylogenomic tree of *E. coli* lineages or other beta barrel porin proteins (Fig. 1A; Supplemental Fig. 1). For instances, MG1655, P12b and UMNK88 are three evolutionarily close strains of lineage A, as was confirmed by *E. coli* core-proteome tree (Fig. 1A). However, the LamB protein sequences from these three strains were located in different phylogenetic clades, while OmpA and OmpC from MG1655/P12b and UMNK88 were also located in different sequence clades (Fig. 1A). Meanwhile, sequences of the strains from different lineages clustered, e.g., LamB proteins of P12b from lineage A, 11368 from lineage B1, and UMN026 from lineage D1 (Fig. 1A). In contrast, PurR, another representative beta barrel porin protein, the sequences from the same *E. coli* lineage were clustered together, and the sequences followed the phylogenetic routes of lineages or core genomes generally (Supplemental Fig. 1).

**Fig 1.**
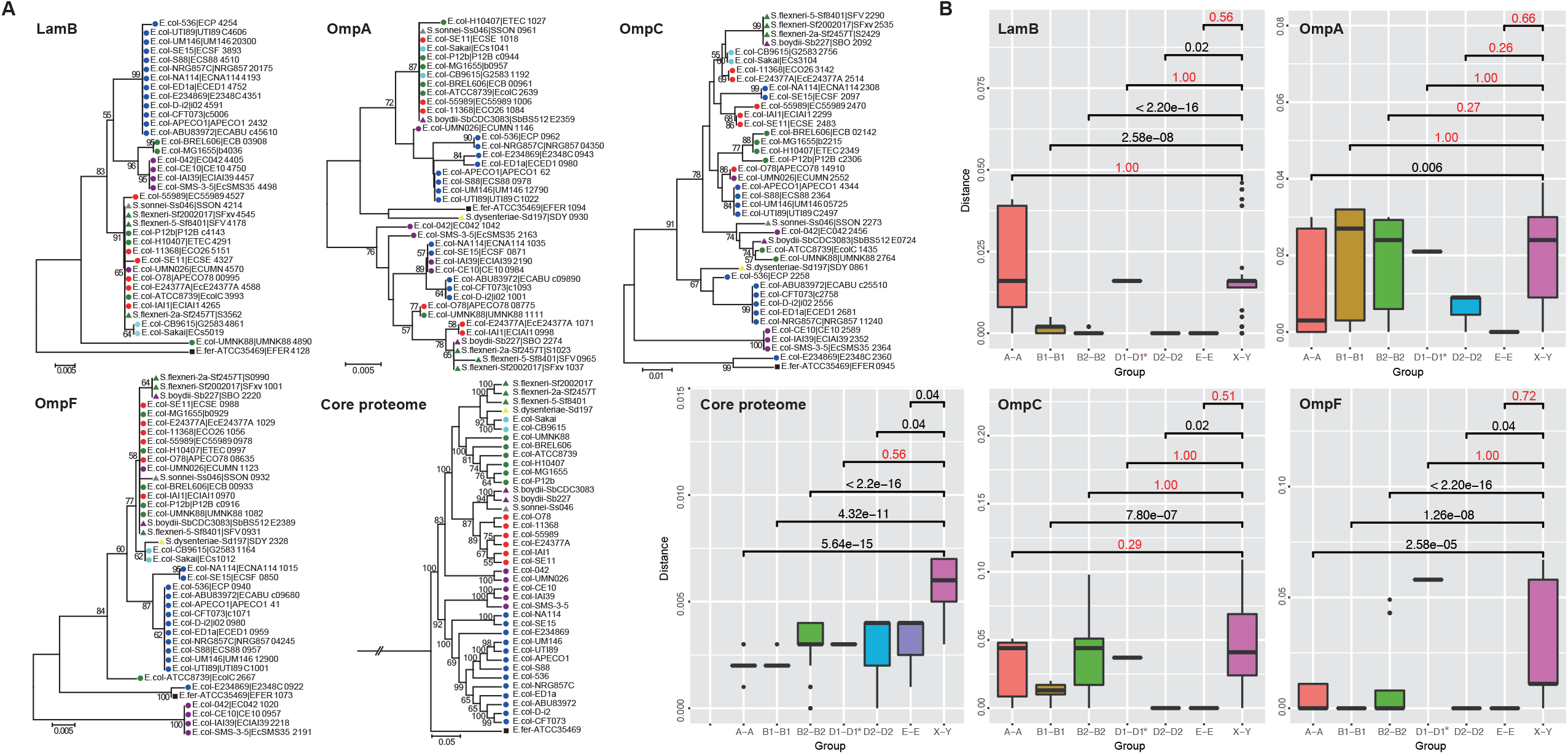
Phylogenetic clustering and genetic exchange testing of *E. coli* porin proteins. **(A)** The Neighbor-Joining un-rooted trees of LamB, OmpA, OmpC and OmpF. For each protein family or the core proteome, the robustness of the phylogenetic tree was examined by bootstrapping tests with 1000 replicates, and the scores were indicated for nodes of no lower than 50 percentages. The gene locus tag for each protein and the strain name were indicated, and strains from the same *E. coli* (or *E. fergusonii*) lineage were labeled with a unique colored sign. **(B)** Genetic exchange testing of porin proteins or the core proteome among *E. coli* lineages. The phylogenetic distance among *E. coli* strains within each lineage was shown and compared with that among strains from different lineages respectively. Mann-Whitney U tests were performed and the *p*-values larger than 0.05 were highlighted in red. ‘A-A’, ‘B1-B1’, ‘B2-B2’, ‘D1-D1’, ‘D2-D2’ or ‘E-E’ represented the strain pairs from the same lineage A, B1, B2, D1, D2 or E respectively. ‘X-Y’ represented pairs of strains from different lineages. The lineage with only two representative strains was indicated with asterisk.

The within-lineage amino acid substitution rates of LamB, OmpA, OmpC and OmpF were compared with inter-lineage substitution rates of the corresponding proteins, respectively (Fig. 1B). The comparison was also performed for the core proteome of *E. coli*, which served as a control. For the control, all the within-lineage rates except D1 were significantly smaller than the inter-lineage rate (Fig. 1B; Mann-Whitney U tests with Bonferroni correction, *p* < 0.05). There were only two representative strains included for lineage D1, as could have led to the lack of significance. For the four porin proteins, the significance was generally lowered, with more than one lineage showing non-significant difference between intra- and inter-lineage phylogenetic distance measured by the substitution rates (Fig. 1B; *p* > 0.05). The results suggested that the mosaic-clustering patterns of the four porin proteins were more likely caused by gene recombination rather than lineage-associated spontaneous mutations. Like *fhuA*, the genomic loci for these genes were conserved, showing large collinearity and synteny among different strains of *E. coli*, suggesting that there were likely within-gene microbial recombination events rather than large-scale horizontal gene transfers that caused the mosaic-clustering patterns observed for the genes (Supplemental Fig. S2).

### 2. Sequence patterns of local recombination hotspots within *E. coli* porin proteins

Multiple sequence alignment showed that most positions of the LamB sequences were conserved for amino acid composition except for a 19-aa region, P406-424 (Fig. 2A). The local highly-variable region (HVR) independently exhibited the clustering patterns exactly like that of the full-length LamB protein sequences (Fig. 2B), while the remaining regions in concatenation formed a phylogenetic tree that is generally consistent with the phylogenomic tree or cannot distinguish different known *E. coli* lineages due to the high sequence conservation (Fig. 2C). The divergence among strains of the same lineage and cross-lineage clustering of the local variable protein sequences further suggested frequent local recombination for *lamB* genes among *E. coli* strains intuitively. According to the phylogenetic clustering results and sequence patterns of the P406-424 region, LamB was classified into 4 groups (LamB1∼4) (Fig. 2A-B). Different groups showed apparent sequence diversity. Within each group, mutations also happened frequently which might caused amino acid changes, and might generate novel, positively-selected patterns that broadened the space of local recombination (e.g., LamB2-1 and LamB2-2; Fig. 2A).

**Fig 2.**
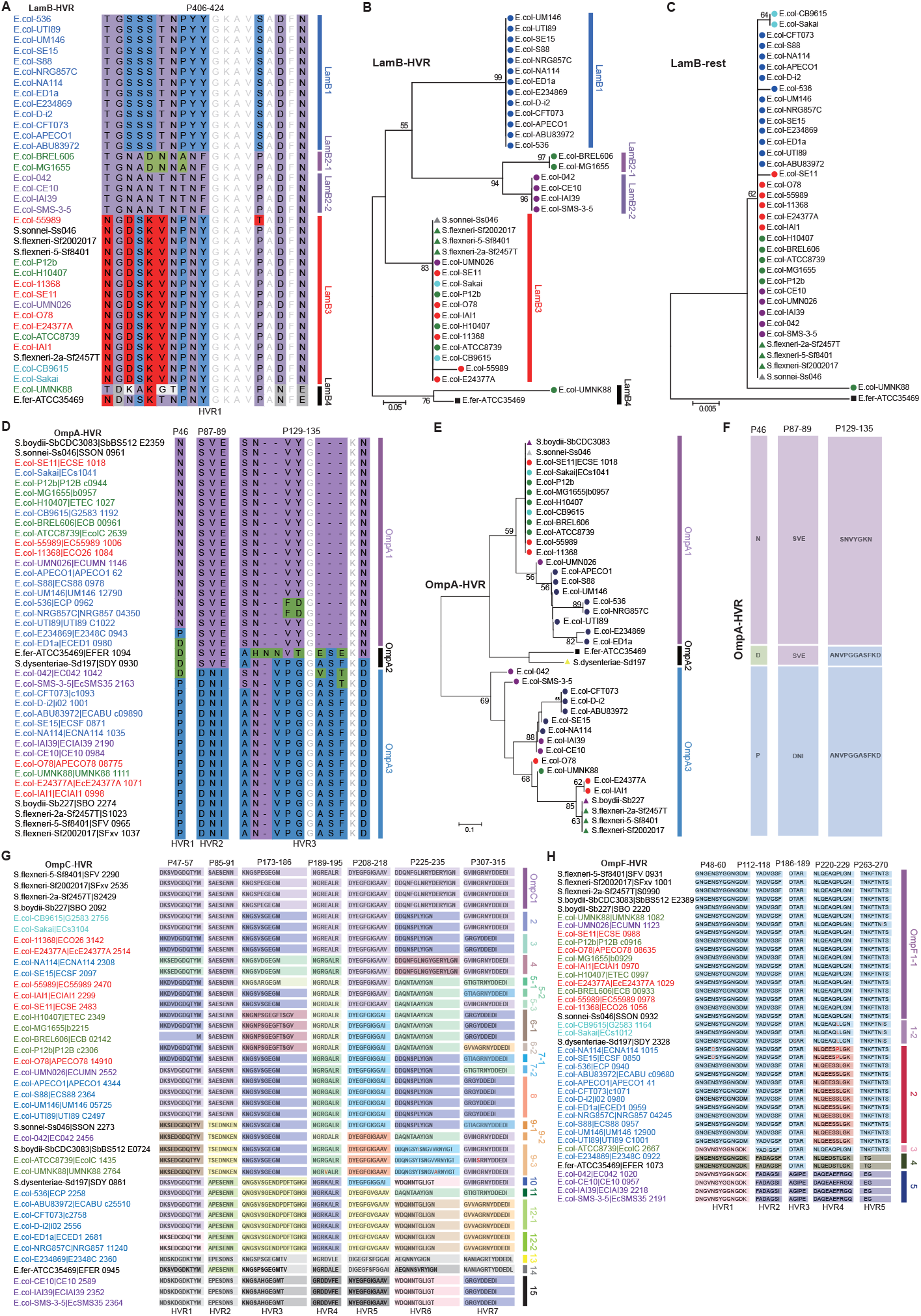
Sequence diversity of highly variable regions within *E. coli* porin proteins. The sequence patterns of highly variable regions (HVRs) of LamB, OmpA, OmpC and OmpF were shown in **(A), (D), (G)** and **(H)** respectively. For OmpC and OmpF, the different patterns for each HVR were shown in different background colors. The Neighbor-Joining tree of the LamB HVR fragment and that of concatenated non-HVRs were shown in **(B)** and **(C)** respectively. The OmpA tree based on the concatenated HVRs was shown in **(E)**. The trees were un-rooted, and bootstrapping tests were performed with 1000 replicates. Strains from the same lineage were labeled with a unique colored sign. The combination of HVR patterns for each OmpA phylogenetic group was shown in **(F)**.

Similar to LamB, OmpA, OmpC and OmpF had also showed the mosaic sequence pattern of conservation for major positions and diversity for local regions (Fig. 2D-H). OmpA, OmpC and OmpF all had multiple HVRs (Fig. 2D, G and H). Similarly, the concatenated HVR sequences of OmpA (or OmpC/F) generated a clustering tree like that derived from full-length protein sequences (Fig. 2E). Different HVRs of OmpA, OmpC and OmpF showed varied sequence patterns, and combined into different patterns, i.e., 3, 15 and 5 major groups for OmpA, OmpC and OmpF, respectively (Fig. 2F-H). HVRs within each group also mutated frequently, and sub-groups could be formed by the mutations or shorter-fragmental recombination events.

### 3. HVRs are located within the outer-membrane convex loops

Tertiary structure of the porin proteins of *E. coli* MG1655 was predicted by different methods. LamB, OmpC and OmpF folded into the classical conformation of beta-barrel outer-membrane spanning proteins, while OmpA showed two separated domains connected by a long flexible loop and one of the domains formed the structure of a beta barrel (Fig. 3A). As observed in FhuA, most HVRs of *E. coli* LamB, OmpA, OmpC and OmpF were located in the extracellular interfaces of the beta barrels, while the remained regions, i.e., transmembrane domains and cytoplasmic interfaces, were quite conserved (Fig. 3A).

**Fig 3.**
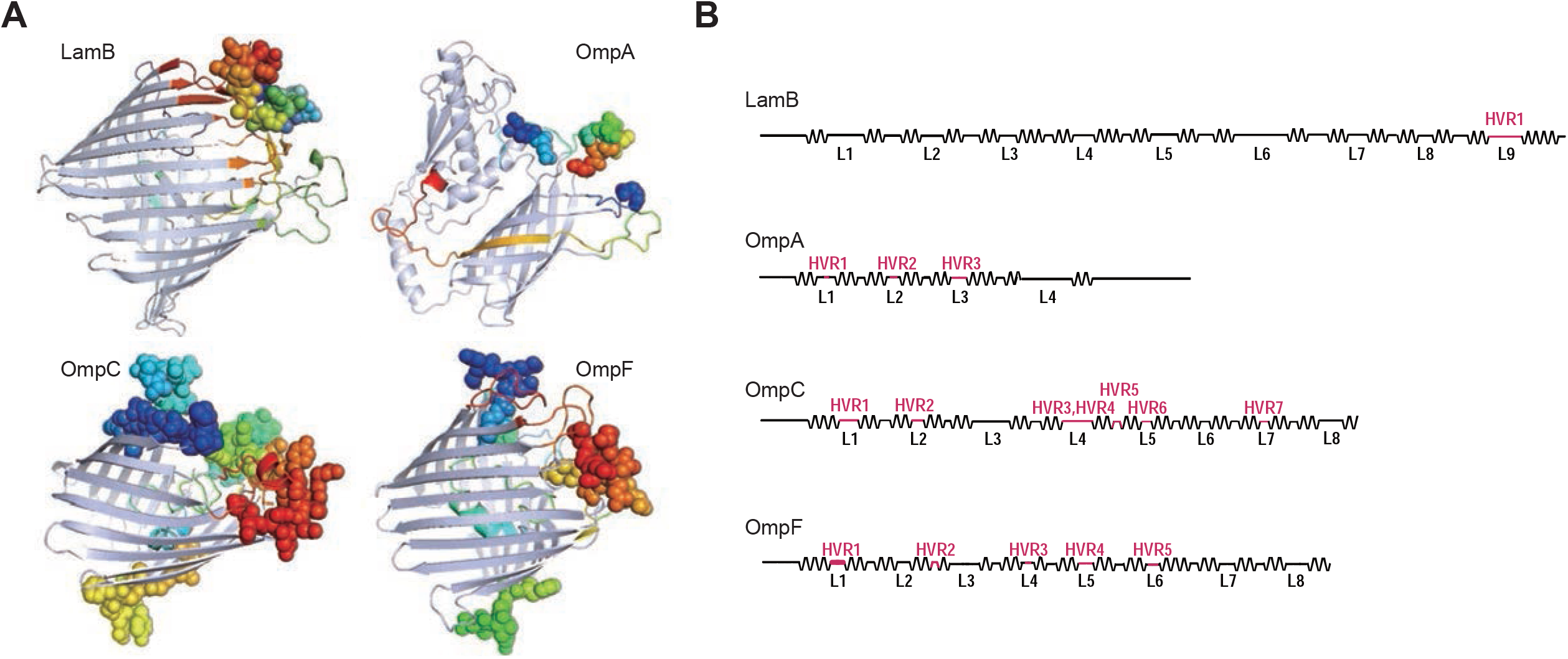
The structural location of HVRs in *E. coli* porin proteins. **(A)** Tertiary structure of MG1655 LamB, OmpA, OmpC and OmpF. The extracellular loops predicted by TMHMM were highlighted in different colors. HVRs were shown in spheres. **(B)** Transmembrane toplogy represented protein sequences of LamB, OmpA, OmpC and OmpF. The topology was predicted with TMHMM. Each extracellular loop was indicated with ‘L’ and the order number. The HVRs were highlighted in red.

Specifically, the HVRs were located within the 9^th^ extracellular loop (L9) for LamB, L1/L2/L3 for OmpA, L1/L2/L4/L5/L7 for OmpC and L1/L3/L4/L5 for OmpF, respectively (Fig. 3B). It was noted that nearly all the HVRs were located at the apex of the convex loops (Fig. 3A). As exceptions, two HVRs, i.e., HVR5 of OmpC and HVR2 of OmpF, were located within the loops within the cytoplasmic interfaces (Fig. 3A-B).

### 4. Structural and functional relevance of the HVRs

The *E. coli* porin proteins of different groups were modeled for the tertiary structure. For LamB, OmpC or OmpF, different HVR groups showed largely homologous global topology (Supplemental Fig. S3). The two big domains of OmpA formed by the N- and C-terminal protein sequences respectively did not show apparent overall difference among the HVR groups either, and yet the distance and relative location between the two major domains varied, which were determined by the angle formed by the long flexible loop connecting the domains (Supplemental Fig. S3).

Different from the general topology of whole proteins, the local structure of extracellular surface formed by the HVRs showed apparent differences for each of the porin families among HVR groups (Fig. 4-7). The structural variations were likely related with protein functions, especially in avoiding bacteriophage binding.

**Fig 4.**
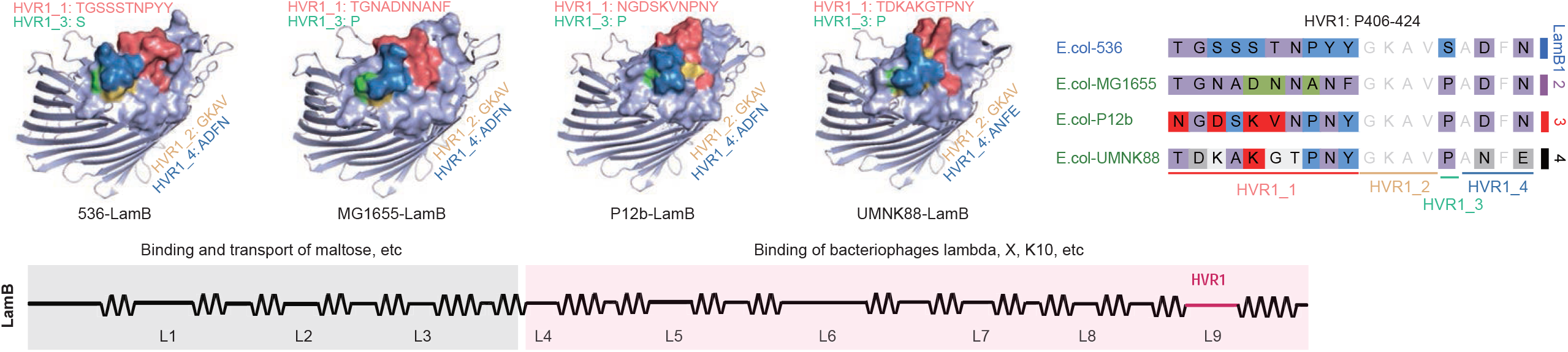
Tertiary structure of *E. coli* LamB protein groups. The sub-regions of HVR1 and their sequences were indicated and highlighted in different colors. The local structure of the sub-regions of HVR1 was shown in colored spheres. The diagram at bottom showed the regional function of LamB.

**Fig 5.**
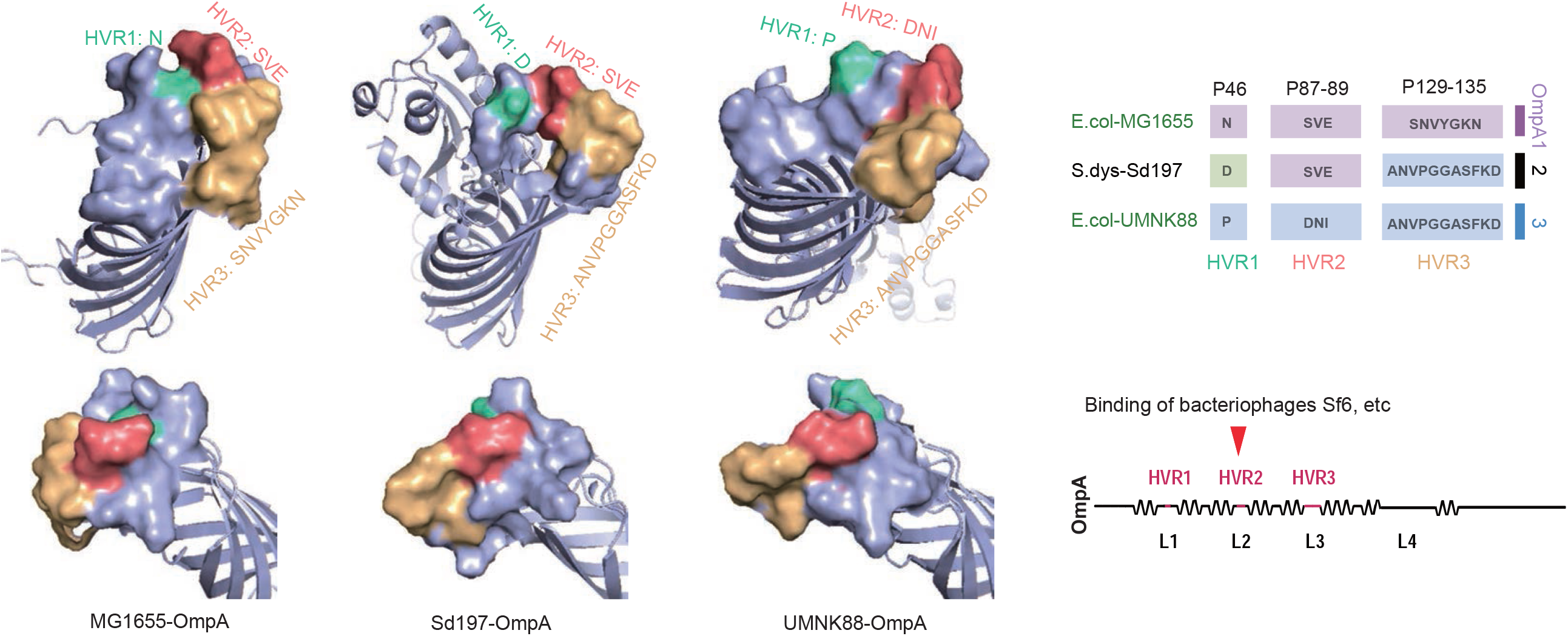
Tertiary structure of *E. coli* OmpA protein groups. The HVRs and their sequences were indicated and highlighted in different colors. The local structure of the HVRs was shown in colored spheres. The known binding sites of bacteriophages were indicated in the diagram.

**Fig 6.**
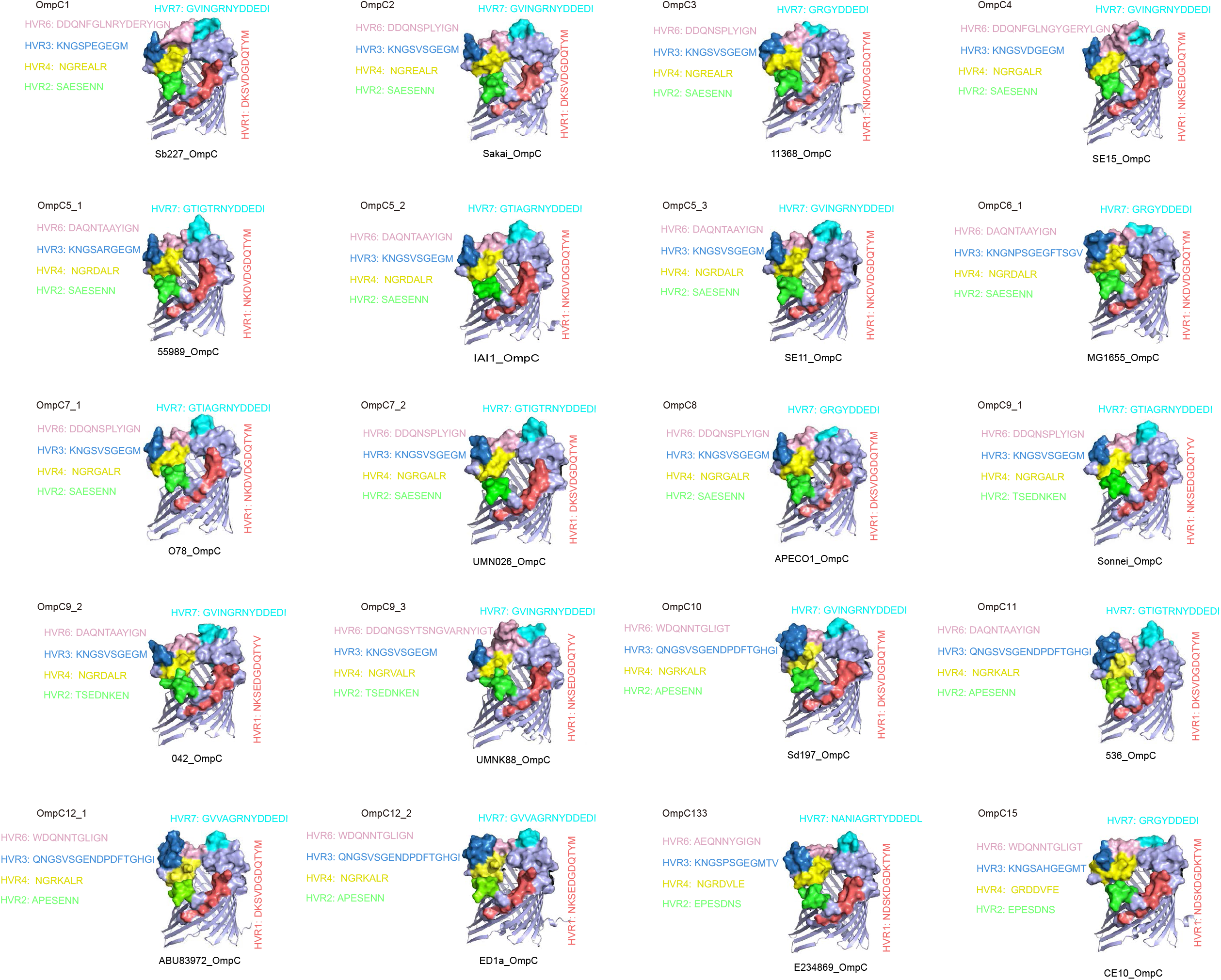
Tertiary structure of *E. coli* OmpC protein groups. The sequences of extracellular HVRs were highlighted in different colors. The local structure of the HVRs was shown in colored spheres.

**Fig 7.**
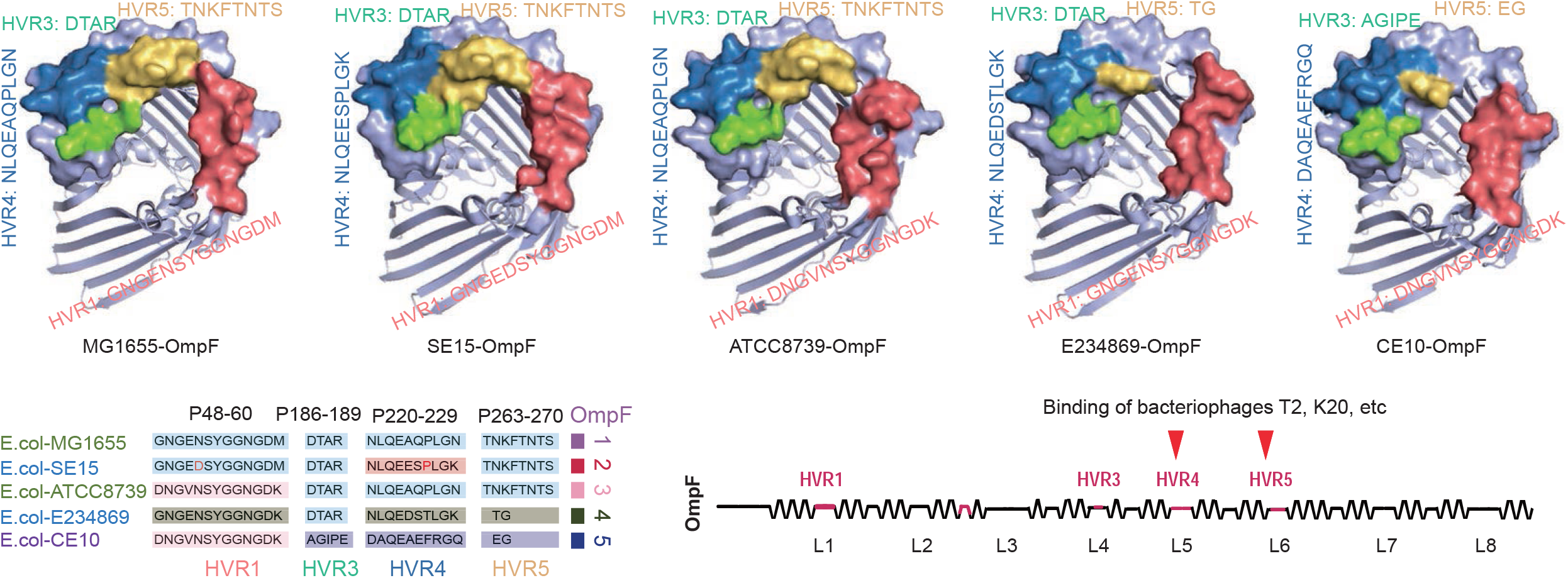
Tertiary structure of *E. coli* OmpF protein groups. The extracellular HVRs and their sequences were indicated and highlighted in different colors. The local structure of the HVRs was shown in colored spheres. The known binding sites of bacteriophages were indicated in the diagram.

In LamB, there was only one HVR (HVR1), which could be divided into 4 sequential sub-regions, HVR1_1 through 4 (Fig. 4). HVR1_1 showed most typical sequence divergence and HVR1_3 was conserved (‘GKAV’) among all the LamB groups. It was noticed that, while the overall interface formed by the whole HVR1 varied, the structure of HVR1_1 and adjacent loci changed most extensively (Fig. 4). LamB is a malto-oligosaccharide-selective pore protein which binds and transports maltose and related bacterial energy source substances. It is also a receptor for the entry of various bacteriophages (Szmelcman and Hofnung 1975; Desaymard et al., 1986; Heine et al., 1987; Chatterjee and Rothenberg 2012). Based on previous reports, the maltose-binding sites were mostly located at the N-terminal ∼1/3 full length of the protein sequence, while the C-terminal amino acids contained key sites that were required by bacteriophages to bind for entry (Desaymard et al., 1986; Heine et al., 1987). HVR1 happened to locate in L9 of the C-terminal part of the protein (Fig. 4), where critical bacteriophage binding residues were reported to locate (Desaymard et al., 1986; Heine et al., 1987). Therefore, the sequence and structure diversity within HVR1 could potentially assist bacteria to avoid the binding and invasion of various phages. No apparent changes were found for other regions or the whole beta-barrel structure, guaranteeing the normal protein function for transport of maltose and other nutrient required for bacterial survival.

OmpA is the receptor of Sf6 and related bacteriophages, while the L2 loop is critical for Sf6 binding (Parent et al., 2014; Porcek and Parent 2015). HVRs 1∼3 of OmpA happen to be located in L1∼3 respectively (Fig. 5), while the bi-residues ‘NI’ in the *Shigella flexneri* HVR2 were confirmed to influence the efficiency of Sf6 entry (Porcek and Parent 2015). HVRs 1∼3 formed an interface with HVR2 in the center flanked by HVR1 and HVR3, diversifying strikingly for the three groups of *E. coli* OmpA (Fig. 5). The sequence of HVRs 1∼3 of *E. coli* UMNK88 OmpA is completely identical to that of *S. flexneri* (Fig. 2B). The HVR2 appeared protruding in UMNK88, but flat and at the same horizontal line with HVR1 and HVR3 in *Shigella dysenteriae* Sd197 where ‘DNI’ was replaced with ‘SVE’ (Fig. 5). In MG1655, the HVR2 became protruding more strikingly because the significant mutations in HVR1 and HVR3 (Fig. 5). The variations in HVR1∼3, especially HVR2, were most likely associated with the interaction with bacteriophages.

The loops L1, L4 and L5 in OmpC, corresponding to HVR1, HVR3 and HVR4, and HVR6 respectively (Fig. 3), were essential for attachment of phage T4, and also maybe for GH-K3 and S16 (Suga et al., 2021; Marti et al., 2013; Cai et al., 2018). L4 (HVR3 and HVR4) is also an important binding site for other OmpC-specific phages, Tu1b, SS4 and Hy2 for instances (Vakharia and Misra 1996). Besides HVR1, HVR3, HVR4 and HVR6, there are other extracellular regions, i.e., HVR2 located in L2 and HVR7 in L7 (Fig. 3), all of which showed large sequence diversity (Fig. 2). Mutations in individual HVRs independently caused microstructure variations, and the combination of diverse sequence patterns in each HVR led to wide and extensive changes of OmpC extracellular interface (Fig. 6).

OmpF is also an important bacteriophage receptor, mainly responsible for the transport of T2-like and other phages. Mutations in the L5 and L6 that coincides with HVR4 and HVR5 of OmpF respectively blocked the binding of phage K20 to E. coli K12 without influence of the OmpF channel activity (Traurig and Misra 1999). Structure modeling also disclosed different appearance for HVR4 and HVR5, especially the latter, among different OmpF groups (Fig. 7). The domain volume and interface formed by OmpF HVR5 varied a lot since there was frequent gain or loss involving multiple residues in the regions, combined with wide substitutions (Fig. 7). In OmpF proteins of *E. coli* MG1655 and SE15, HVR4 and HVR5 were contiguous, while the HVRs were separated with a cleft in ATCC8739 and they were separated far away in E234869 and CE10 (Fig. 7). The conformation changes of both individual HVRs and the whole interface could be associated with the binding and entry efficiency of bacteriophages.

Taken together, the results demonstrated that all the four porin proteins showed apparent structure changes at the HVR regions but without global or channel topology difference. Most of the HVRs were reported to contain critical recognition sites for various bacteriophages, and therefore the diversity in HVRs could probably endue the bacterial strains resistance from phage invasion. However, the conservation of global structure, cytoplasmic interfaces and other extracellular surfaces ensured the maintenance of the transporter and binding activity for substances useful for bacterial survival.

## Discussion

Previously, we reported a gene encoding a siderphore transporter, *fhuA*, which showed an extraordinary evolutionary pattern, i.e., with combination of negative selection, positive selection and apparently local recombination simultaneously (Wang et al., 2018). The kind of within-molecule mosaic evolution is hardly detected with established tools, and therefore *fhuA* and other genes were only reported under positive selection but no recombination was identified (Chen et al., 2006; Petersen et al., 2007). In this research, we broadened observations to other beta-barrel porin proteins that were also reported to be under positive selection (Chen et al., 2006; Petersen et al., 2007). Interestingly, at least 4 additional proteins were found following the similar evolutionary routes. Others not included in the report did not necessarily show traditional evolutionary patterns, but were excluded since their local-recombination patterns were not as typical as FhuA or the other four porin proteins. Despite no direct evidence and the possible structure-constrained co-convergence mutations (Hu et al., 2016), the locally fragmental convergence patterns within proteins of the strains from different lineages supported the most likely local sequence recombination events. Such mosaic evolution patterns have also been identified within multiple porins in other species, e.g., OprD in *Pseudomonas* (Chevalier et al., 2007), PorA and PorB in *Neisseria* (Derrick et al., 1999; Posada et al., 2000; Urwin et al., 2002) OmpL1 in *Leptospira* (Haake et al., 2004), OmpA in *Chlamydia* and *Wolbachia* (Millman et al., 2001; Baldo et al., 2010) and OmpF in *Yersinia* (Stenkova et al., 2011). Besides, mosaic genetic exchange events were also identified from bacterial adhesins and other outer membrane beta-barrel proteins (Falush et al., 2001; Nell et al., 2014; Kumar et al., 2018). The mosaic evolution could represent a common route undergone by bacterial outer membrane proteins.

The proteins studied in this study or elsewhere under mosaic evolution are mainly multi-functional outer membrane proteins. Typically, they have important functions, for example, import of nutrient or energy source substances essential for bacterial survival (Szmelcman and Hofnung 1975; Hantke and Braun 1975; Millman and Dean 2001). Phages, virulence factors such as colicins and microcins deployed by competitors, antibiotics or antibodies can also hijack these proteins to enter and kill the recipient bacteria (Braun 2009; Chatterjee and Rothenberg 2012; Baker and Casjens 2014). The receptors must hide themselves from recognition by the harmful factors but maintain the capability of being recognized and bound by the essential substances and the transport activity. Therefore, from the evolutionary traces, within these proteins, the fragments involved in transport activity and the binding sites of substances essential for bacterial survival should be conserved while the recognition sites of disfavored factors should be under positive selection and mutate extensively (Desaymard et al., 1986; Stenkova et al., 2011; Wang et al., 2018). Consistent with the hypothesis, all the porin proteins studied in this study showed globally conserved sequence and structure interspersed with small HVRs with extensive sequential and structural diversity. Within these proteins, nearly all the reported critical phage-binding sites coincided with the mosaic HVRs, and the known recognition sites of bacteria-needed substrates or regions involved in transport activity were seldom located in the HVRs or showed structural variations. The type of mosaic evolution has also been identified from multi-cellular eukaryotic organisms although different evolutionary mechanisms could be involved. For example, there are similar multi-functional receptor proteins usurped by fatal viruses in human or other animals, e.g., ACE2 bound by SARS-CoV-2 related viruses, the co-receptor of HIV-1 and *Yersinia pestis* CCR5, etc., and local diversities with functional relevance have been identified from these proteins (Guo et al., 2020; Mummidi et al., 1998).

Despite the observations of local recombination in the porin proteins, it remains unclear why the bacteria adopt the way rather than positive Darwinia selection predominantly mediated by spontaneous mutations. Both recombination and mutation based positive selection should happen within HVRs of the bacterial proteins and play important roles in the evolutionary processes. Genetic recombination is common within bacteria in natural environment and happens frequently (Guttman and Dykhuizen 1994; Vos and Didelot 2009). When a bacterial population encounters the attack of phages or other harmful substances, both genetic recombination and spontaneous mutations could happen that help the population survive from the disfavored environment. However, the positive selection mediated by random mutations lasts for a long period, involving a slow fitting process through continuous gene mutations for generations. If there are polymorphic gene fragments from other bacteria that have the capability to resist from the invasion of the encountered phages or other damaging factors and have experienced the long selective and fitting process, and if homologous recombination in individual bacteria involving the gene or local fragments happens, such strains could probably have the fitting advantages over the positively selected ones mediated by random mutations (Vos 2009). It explains the observations of predominant local recombination in the proteins. However, mutations and their mediated positive selection continue for better adaptation of the bacterial population lives in the environment. Meanwhile, the spontaneous mutations create new polymorphic forms of gene sequences, further enlarging the pool for recombination.

It should be noted that the structure for the different groups of porin proteins was predicted with different state-of-the-art methods, including a classical homology-modeling based method and another ab-initio one proposed most recently that was reported with best performance at present (Kelley et al., 2015; Baek et al., 2021). Except for the angles between the two major domains in OmpA formed by the long flexible coil, the global structure for each porin family was conserved, and consistent between the prediction results of different prediction methods. Domain and local structure of the proteins predicted by the two methods were generally consistent and therefore reliable. Similar to the known sites critical for the entry of known phages, in the study, we identified more and longer HVRs in most of the porin proteins, which also showed extensive sequence and local structure changes. Meanwhile, there could be a phage repertoire larger than what we have known, that takes the porin proteins as receptors for entry into host cells. Other unknown toxins, antibiotics or immune surveillance could also target these porins. Therefore, the porins under mosaic evolution and the HVRs identified in the study could facilitate the studies on new phage-bacteria co-evolutionary mechanisms, identification of their possible critical binding sites, and development of new anti-bacterial drugs or treatment regimens.

## Materials and Methods

### 1. Bacterial strains, genomes and protein sequences

The genes reported under positive selection in *E. coli* were noted down from previous studies and merged (Chen et al., 2006; Petersen et al., 2007), and the protein sequences from the model strain MG1655 were downloaded from NCBI Protein database (https://www.ncbi.nlm.nih.gov/protein). Representative strains for major *E. coli* lineages, their lineage information, genome sequences, genome annotation files, genome-derived proteome sequences, core genome and its derived proteome were retrieved from our previous study (Wang et al., 2018). A representative strain of *E. fergusonii*, ATCC35469, was used as outgroup or control and its genome or proteome information was also downloaded. As before, *Shigella* species were considered as phylogenetic sub-branches of *E. coli* (Wang et al., 2018).

### 2. Phylogenetic analysis

Phylogenetic clustering was performed to the full-length porin proteins or peptide fragments. Amino acid sequences were aligned with ClustalW (https://myhits.sib.swiss/cgi-bin/clustalw) using the default parameters. Phylogenetic trees were built with both Neighbor-Joining (NJ) and Maximum Likelihood (ML) methods implemented in MEGA 6.0 (Tamura et al., 2013), while only the NJ trees were illustrated in the study because the NJ and ML trees appeared consistent between each other for all the studied proteins. Bootstrapping tests were performed for 1000 replicates to assess the robustness of each node of a tree, at which only if ≥50 percentages of the subtrees show consistence the branch could be considered stable. The core-proteome trees of *E. coli* were re-built according to the methods described previously (Wang et al., 2018).

### 3. Genetic exchange testing

To assist screening the ingenic local genetic exchange events, a simplified hypothesis based testing was proposed. *H*_*0*_: No local genetic exchange happened among strains from different lineages; *H*_*1*_: Local genetic exchanges happened among strains from different lineages. Suppose *H*_*0*_ holds. The phylogenetic distance among strains from the same lineage should be significantly smaller than the distance among the strains from different lineages, as could be proved in the first place. However, if the intra-lineage distance is larger than or not significantly different from the inter-lineage distance, the starting hypothesis (*H*_*0*_) would be rejected and the optional hypothesis (*H*_*1*_) is accepted. In the study, phylogenetic distance was measured as the adjusted substitution rate of amino acids. The *E. coli* core-genome derived proteome was used as control to prove the smaller intra-lineage distance than inter-lineage distance. Representative *E. coli* strains were selected, followed by calculation of the phylogenetic distance for porin proteins or core proteome between each pair of strains of the same lineage (intra-lineage) or from different lineages (inter-lineage). Mann-Whitney U tests were performed to compare the difference between intra-lineage and inter-lineage distance. Bonferroni corrections were performed to the *p*-values, since multi-testing was involved. For all the statistic testing, the threshold of type I error was preset as 0.05.

### 4. Structure modeling

The transmembrane topology of LamB, OmpA, OmpC and OmpF was predicted with a Hidden Markov Model (HMM) based method, TMHMM 2.0 (Krogh et al., 2001). The tertiary structure was predicted with Phyre2 by homology modeling (Kelley et al., 2015) and RoseTTAFold *ab intio* (Baek et al., 2021). Besides the predicted accuracy or resolution estimated by the tools themselves, the results predicted with the two methods for each protein were compared between each other and the consistence was also based to assess the reliability. For each protein, the coordinates of atoms for each amino acid were recorded in PDB format, and the 3D structure was illustrated with PyMOL (https://pymol.org/2/).

## Acknowledgements

The study was supported by a Natural Science Fund of Shenzhen (JCYJ20190808165205582) to Y.W. Z.C. was supported by a Fund for the Cultivation of Guangdong College Students’ Scientific and Technological Innovation, Climbing Program (pdjh2021b0432). X.C. was supported by an Undergraduate Training Program for Innovation and Entrepreneurship of Shenzhen University (no.127) and an Undergraduate Science and Technology Innovation Program of Shenzhen University.

## Authors’ Contribution

Y.W., Y.C., A.P.W, K.W. and G.Z. conceived the project; X.C. and Z.C. collected the data; X.C., X.C., Z.C., J.W. and Y.W. performed the analysis; G.H. and Q.L. participated in the discussion; X.C., X.C., Z.C., G.H., Q.L. and Y.W. wrote the first draft; A.P.W., K.W., G.Z., and Y.W. revised the manuscript. All the authors approved the final version of manuscript.

## Supplemental Materials

**Fig. S1. The Neighbor-Joining tree of E. coli PurR**. *E. fergusionii* ATCC35469 was used as outgroup. The bootstrapping testing scores were indicated for nodes of no lower than 50. The gene locus tag for each protein and the strain name were indicated, and strains from the same *E. coli* (or *E. fergusonii*) lineage were labeled with a unique colored sign.

**Fig. S2. The synteny of *E. coli* porin genes**. The phylogenetic tree built with *E. coli* core genome derived proteome was shown in the left, and for each strain the array of the porin genes and their flanking genes were shown in the right accordingly.

**Fig. S3. Tertiary structure of various protein groups for LamB, OmpA, OmpC and OmpF**. Each protein was a representative of a group defined by the HVR patterns and combinations. The structure was predicted with RoseTTAFold and Phyre2, and compared, while only the structure predicted by RoseTTAFold was shown.

